# A guide to the manufacture of sustainable, ready to use in vitro platforms for the electric-field pacing of cellularised 3D porous scaffolds

**DOI:** 10.1101/2022.10.04.510868

**Authors:** Matteo Solazzo, Michael G. Monaghan

## Abstract

Electrical activity is a key feature of most native tissues, with the most notable examples being the nervous and the cardiac systems. Modern medicine has moved towards the mimicking and regenerations of such systems both with in vitro models and therapies. Although researchers have now an increased repertoire of cell types and bio-physical cues to generate increasingly complex in vitro models, the inclusion of novel biomaterials in such systems has been negligible, with most approaches relying on scaffold-free self-assembling strategies. However, the rapid development of functional biomaterials and fabrication technologies - such as electroconductive scaffolds – warrants consideration and inclusion of materials, with recent evidence supporting the benefit of incorporating electrically active materials and their influence on the maturation of cardiac cells and tissues. In order to be manipulated in bioreactor systems, scaffold-based in vitro models require bespoke rig and bioreactors that vary from those commonly used for scaffold-free systems. In this work, we detail methods to rapid prototype an electrical pacing bioreactor and R3S - a Rig for Stimulation of Sponge-like Scaffolds. As a proof of concept and validation we demonstrate that these systems are compatible with isotropic and anisotropic porous scaffolds composed of collagen or poly(*3,4*-ethylene dioxythiophene):polystyrene sulfonate (PEDOT:PSS). External pacing of C3H10 cells on anisotropic porous scaffolds led to a metabolic increase and enhanced cell alignment. This setup has been designed for pacing and simultaneously live tracking of in vitro models. This platform has wide suitability for the study of electrical pacing of cellularized scaffolds in 3D in vitro cultures.

## 1. Introduction

Bioreactors are controllable dynamic systems used to recapitulate biophysical stimuli, scale up cultures in vitro to generate sufficient numbers of therapeutic cells, growth factors, proteins and hormones, and study physiological responses ex vivo ^1^. Such stimuli include strain and stress, pressure, flow, and the subject focus of this paper: electrical pacing. Pacing is a method commonly used with engineered heart tissues derived from primary cardiomyocytes and cardiomyocytes derived from pluripotent stem cell sources ^2^, neural applications in the case of neuronal cultures ^3^, and skeletal muscle cultures ^4^. More recently, the use of electrical pacing has been explored in non-traditional tissues such as skin ^5^, bone ^6^, and immune cells ^7^.

Many electric field-based systems investigate cells embedded in fibrin ^8^ or collagen ^2^ gels, in the form of tubules ^8^, rings ^9^ or patches ^10^. Engineered tissues are also often generated using biologically-derived hydrogels, typically based on fibrin or collagen, enabling spontaneous self-assembly, and has been applied to the generation of engineered heart tissues since 1997 ^11^. For example, engineered heart tissues are often incorporated into bioreactors capable of delivering diverse cues (e.g. shear rate, mechanical constrain, electrical stimulation), and these stimuli have been shown to support the maturation of induced pluripotent derived cardiomyocyte (iPSC-CMs) cells towards more adult-like phenotypes with functionality ^12^. In particular, electrical pacing can synchronise cell beating with an effect similar to the action potential generated in the sinoatrial node, and it can also be used to tune the beating frequency of the engineered heart tissues itself ^13–14^ and study inherited genetic defects in engineered heart tissues ^15^.

A less explored strategy in engineered tissues subjected to electric field stimulation is seeding cells onto a pre-existing preformed scaffold, possessing a predefined geometry and stiffness ^16^. Due to simplicity of fabrication, many researchers have focussed on the scaffold-free self-assembling method, most recently with the generation of a human-size engineered heart tissues ^10^. A possible reason for the lack of scaffold-based strategies is the repertoire of materials and constructs available such as collagen sponges ^17^ or polylactic acid fibrous scaffolds ^16^.

The burgeoning progress of electroconductive biomaterials, especially conjugated polymers (CPs), may impact this trend and rationalise scaffold use for engineered tissues in vitro. These materials have demonstrated potential to assist the regeneration of the heart wall after MI ^18–20^, and it is hypothesised that such electroconductive platforms could provide enhanced electrophysiological features when compared to biologically derived materials ^21^. While their use as 2D substrates in cardiac tissue engineering is gathering momentum ^22^, the inclusion of electroconductive materials into engineered tissues has yet to be fully exploited. The use of porous scaffolds with predefined microarchitecture could directly influence cell response, in particular their spatial orientation. In the case of a highly anisotropic tissues such as myocardium, muscle or nerve fibres, microarchitecture can guide the alignment of cells along the direction of contraction and their fusion into structures ^23–24^. Material stiffness is another physical property known to play a significant role, in promoting the differentiation of progenitor cells towards a desired progeny ^25^.

In this work, we describe the development of an in-house electric field bioreactor system for use with prefabricated porous scaffolds for 3D electric-field stimulation. We apply electroconductive scaffolds based on crystallised poly(*3,4*-ethylene dioxythiophene):polystyrene sulfonate crosslinked with (3-Glycidyloxypropyl)trimethoxysilane (PEDOT:PSS-GOPS) as 3D substrates for in vitro models based on our extensive experience with this material. This in-house bioreactor is capable of applying electrical pacing to monolayer cell cultures as well as to 3D constructs and its construction and design is documented in detail. Furthermore, we conceptualised and manufactured R3S (Rig for the Stimulation of Sponge-like Scaffolds); an in vitro platform designed to maintain scaffolds suspended in media in the same location, while enabling free-contraction of the cell laden constructs and simultaneously providing optical accessibility for both standard and inverted microscopy. This is achieved using custom-made grips incorporated into this platform. We report validation of this system using a C3H10 cell line and demonstrate that scaffold morphology and electrical pacing play a synergistic role on the proliferation and metabolism of cells. Moreover, C3H10 cells preferentially aligned along the direction of the electric field even in the absence of an anisotropic substrate.

Here we provide detailed methods on the fabrication and assembly of this bioreactor system. We expect the information presented in this manuscript will provide researchers the ability to develop their own electric-field bioreactor assembly system to suit a diverse number scaffold-based cultures and various other potential applications.

## 2. Experimental Section

### 2.1. Modelling and 3D printing

Throughout the project, 3D models were designed and then fabricated via rapid prototyping. All software design was performed with Solidworks while 3D printing was achieved with either Original Prusa i3 MK3S (Prusa Research), Ultimaker 3 (Ultimaker BV) or stereolithography (SLA) printer Form 3 (Formlabs). These files are available in the supplementary section.

### 2.2. Processing of 3D sponge-like scaffolds

Porous sponge-like scaffolds were generated using either collagen type I slurry or PEDOT:PSS-based dispersion; with collagen type I as the natural extracellular matrix based sponge material and PEDOT:PSS scaffolds as the electroconductive control which we have worked with extensively in another study ^26^.

Collagen type I was isolated from porcine tendons following as described previously ^26^. The isolated collagen type I was then solubilised in 0.1 M acetic acid to a final concentration of 15 mg/ml. PEDOT:PSS 1.3 wt.% dispersion in water (Sigma-Aldrich, Ireland) was mixed with 3 v/v% GOPS (average M_n_ 236.34, Sigma-Aldrich, Ireland). PEDOT:PSS blends were vortexed for 30 seconds, sonicated for 30 minutes and filtered with a 0.45 μm PVDF syringe filter to remove aggregates.

Porous 3D sponge-like scaffolds were created through lyophilisation using a freeze-dryer (FreeZone, Labconco Corporation, USA). Isotropic scaffolds were produced using standard cell culture multi-well plates, while a custom-made mould containing a bottom stainless-steel layer and a top PDMS (SYLGARD® 184, Farnell, Ireland) was fabricated to induce ice crystal alignment by virtue of the different thermal conductivities of the two materials ^27^. Freezing was performed at −40 °C, for 1 hour after which the temperature raised to −10 °C at a rate of 1 °C per minute, held for 18 hours at a vacuumed environment of 0.2 mbar, ramped to 20 °C at a rate of 1 °C per minute, and finally held for 2 hours.

PEDOT:PSS-GOPS scaffolds underwent annealing treatment in a vacuum oven at 140 °C for 1 hour and were crystallised as previously reported ^26^. Briefly, PEDOT:PSS-GOPS samples - with both isotropic and anisotropic architecture - were submerged into a bath of 100% sulphuric acid for 15 minutes, then washed multiple times using deionised water.

After freeze-drying, collagen scaffolds were crosslinked with *N*-(*3*-Dimethylaminoproypl)-*N*-ethylcarbodiimide hydrochloride (EDC) (E1769, Sigma-Aldrich) and *N*-Hydroxysuccinimide (NHS) (130672, Sigma-Aldrich). A 5:5:1 molar ratio of EDC:NHS:carboxyl groups within the collagen scaffolds was added to pure ethanol and the pH adjusted to between 5.3 and 5.5 ^28^. Crosslinking took place under rocking for 18 hours; samples were then rehydrated and lyophilised one more time.

Scaffolds were finally sectioned at 1 mm thickness with a vibratome (VT 1200S, Leica. Germany). Regular rectangular strips 9 mm long and 2 mm wide were manually cut using razor blade, followed by sterilisation and preconditioning in standard growth media.

### 2.3. Fabrication of pacing bioreactor

To electrically pace cells upon scaffolds, electrical stimulation was delivered via a custom-made bioreactor. We conceptualised and manufactured the system with the use of inexpensive rapid prototyping, gathering inspiration from existing models ^12, 29^. A schematic representation of the setup is shown in **Figure 1**. A set of twelve 22 mm long cylindrical carbon electrodes were cut from a 3 mm large carbon rod (Goodfellow Cambridge Ltd, UK) and a 1 mm large hole was drilled at one end, through which a platinum-iridium wire (Advent Research Materials Ltd, UK) was tied. A mould, consisting of a top and a bottom component (**Figure S1**), was 3D printed to allocate twelve electrodes and, once the two components were sealed together, PDMS Sylgard™ 184 was introduced in the mould. After polymerisation, the mould was opened, and the electrodes could be collected (**Figure 1.A** and **Video 1**). The silicone closures at both extremities of each carbon bar allowed permanent securing of the platinum-iridium wire and avoidance of any liquid infiltration, while also providing an anchorage system to the *lid adapter*. This 3D printed component interfaced between the electrodes and the plate lid (**Figure 1.Bi**, **S2 and Video 2**); it holds 6 pairs of carbon rod electrodes kept at 15 mm distance between each other and ultimately able to pace a 6-well plate. This component was designed to have slots for six screws to secure it to a 6-well plate lid, with openings in the area in between the electrodes for optical accessibility with both upright and inverted microscopes. A 5 mm large space between the interface and the lid was accounted in for the wiring. Electrodes are connected with copper wire to have a configuration in parallel as represented in **Figure S3**. The source ground and positive wires connected externally to a bread board where a waveform generator (RSDG2000X, Radionics) and an oscilloscope (RSDS 1052 DL +, Radionics) were also connected. The oscilloscope serves to validate the signal being generated by the waveform generator.

**Figure 1.**
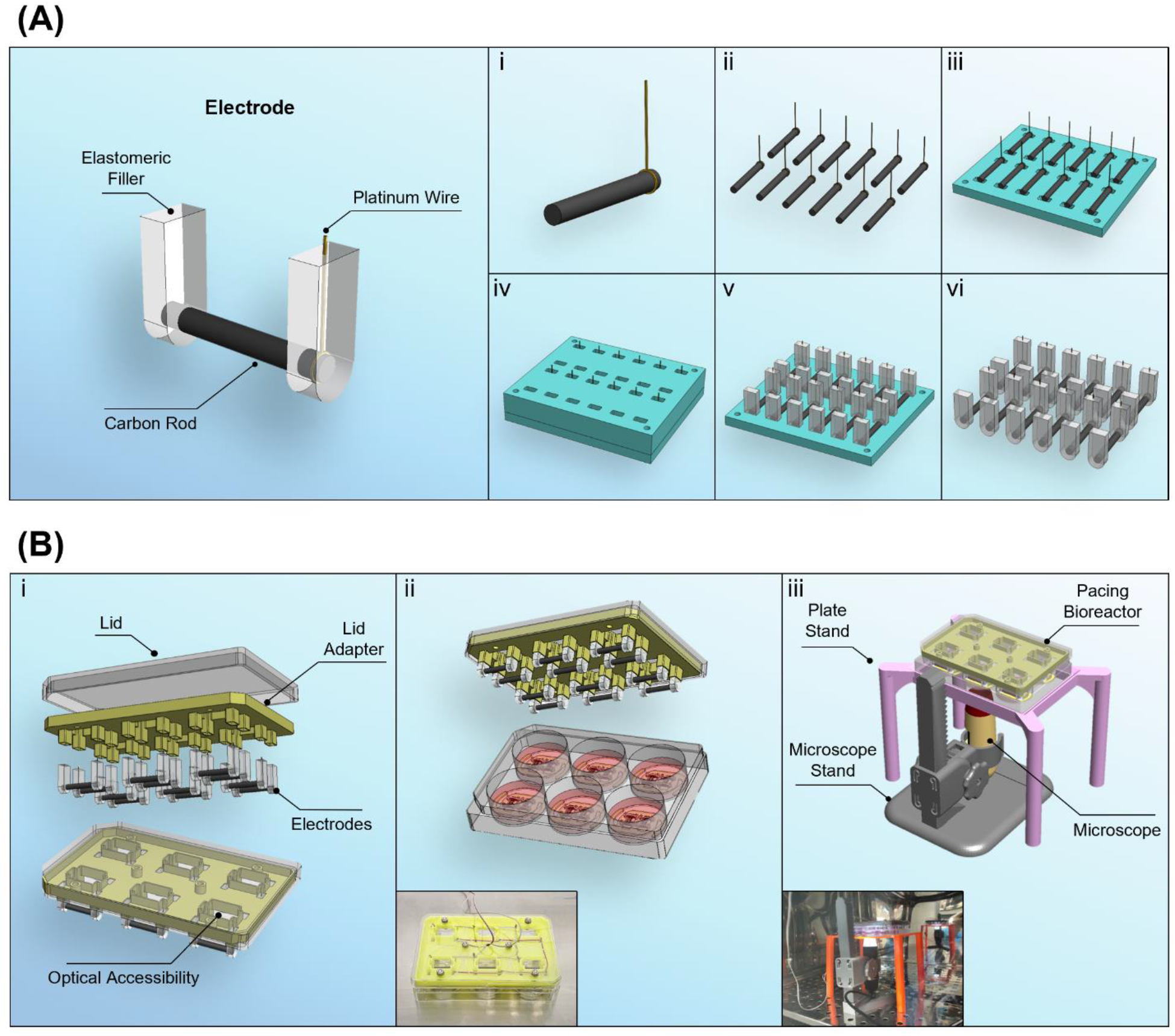
Design and manufacturing of the electrical pacing bioreactor. **(A)** Schematic representation of a carbon electrode. (i-vi) Fabrication sequence of the electrodes. (i) A platinum-iridium wire is tightened to one extremity of a carbon bar. (ii) Step i is repeated until achievement of 12 elements. (iii) Bars are inserted into the bottom component of the mould. (iv) The top component of the mould is secured and pristine PDMS is poured into each hole. (v) Polymerisation is completed and the top component of the mould can be removed. (vi) Once the bottom component is removed, 12 carbon electrodes are achieved. **(B)** Overview of the electrical pacing bioreactor. (i) Exploded and collapsed views of the individual components of the electrical pacing bioreactor. (ii) Schematic and picture of the custom-made cell culture pacing setup. (iii) Schematic and picture of the custom-made setup for the observation and monitoring of scaffolds.

### 2.4. Fabrication and tensile testing of elastomeric bars

Elastomeric bars were designed in dumbbell-like shape and were placed in pairs at each extremity of the scaffold in a sandwich-like fashion. These bars were casted within a multi-component custom-made mould (**Figure 2.Ai-iii**) and had a square cross section with 1 mm long sides (**Figure 2.Aiv**). PDMS blended as a 1:5 ratio of Sylgard 184 and Sylgard 527 was used, to provide a material that could be strong enough to keep the scaffolds in place, but with appropriate flexibility to be deflected by contracting scaffolds.

**Figure 2.**
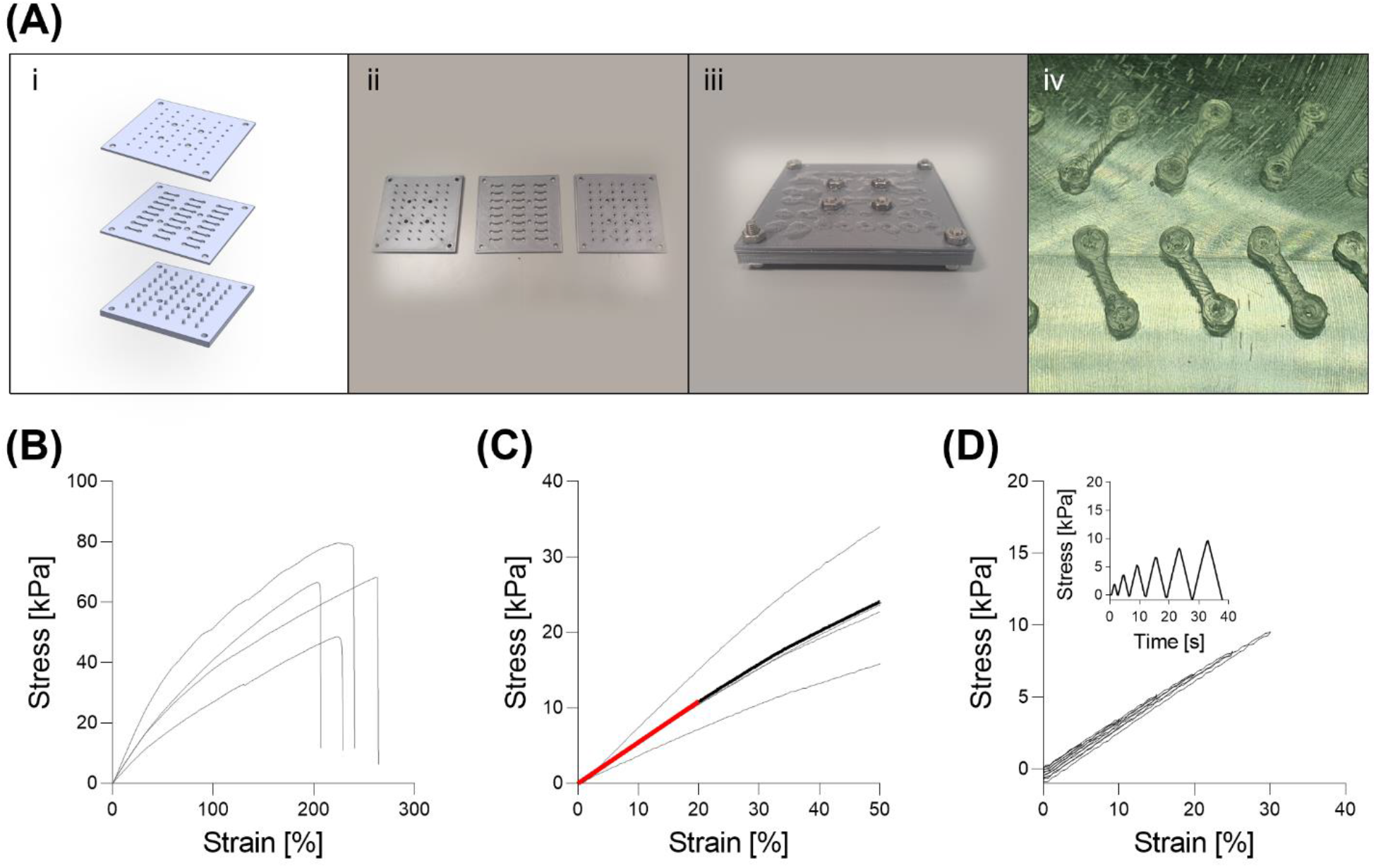
**(A)** Manufacturing of PDMS elastomeric bars. (i) Illustrations of bottom, middle and top components of the mould. (ii) 3D printed components of the mould. (iii) Assembled Mould that has been filled with pristine PDMS. (iv) A set of manufactured PDMS elastomeric bars. **(B-D)** Mechanical characterisation of PDMS bars via uniaxial tensile test. **(B)** Tension until rupture. **(C)** Tension in the range 0-50% strain with mean linear interpolation in red. **(D)** Stress-strain and stress-time curves representing a standard response within cycles from 5% to 30% strain.

Uniaxial tensile tests were performed on elastomeric bars to characterise their stiffness. First, a cyclic sequence with increasing strain from 5% to 30% by 5% increments was performed to assess the absence of hysteresis in dynamic conditions. Then, a ramp until failure followed to verify the rupture limit (**Figure 2**). Bars had a mean elastic modulus of 54.3 ± 15.7 kPa and no evidence of hysteresis was observed in the tested range.

### 2.5. R3S: Rig for the Stimulation of Sponge-like Scaffolds

For the stimulation and optical monitoring of 3D sponge-like scaffolds, a specific rig that could maintain scaffolds in a stable position was developed. In addition, for the application of such a system to tissue engineering, this rig should not impede the typical contraction of the scaffolds generated by beating skeletal muscle myocytes or cardiomyocytes. The system presented in **Figures 3.A** is constituted by a main body and four elastomeric bars. The main body (**Figure S4 and Video 3**) was designed to accommodate two 3D scaffolds that were to be kept parallel with the electric-field generated by the bioreactor, that could be observed from optically accessible chambers placed underneath. On the side and at the extremities of each of these chambers, a pillar with a sharp tip anchors the elastomeric bars.

**Figure 3.**
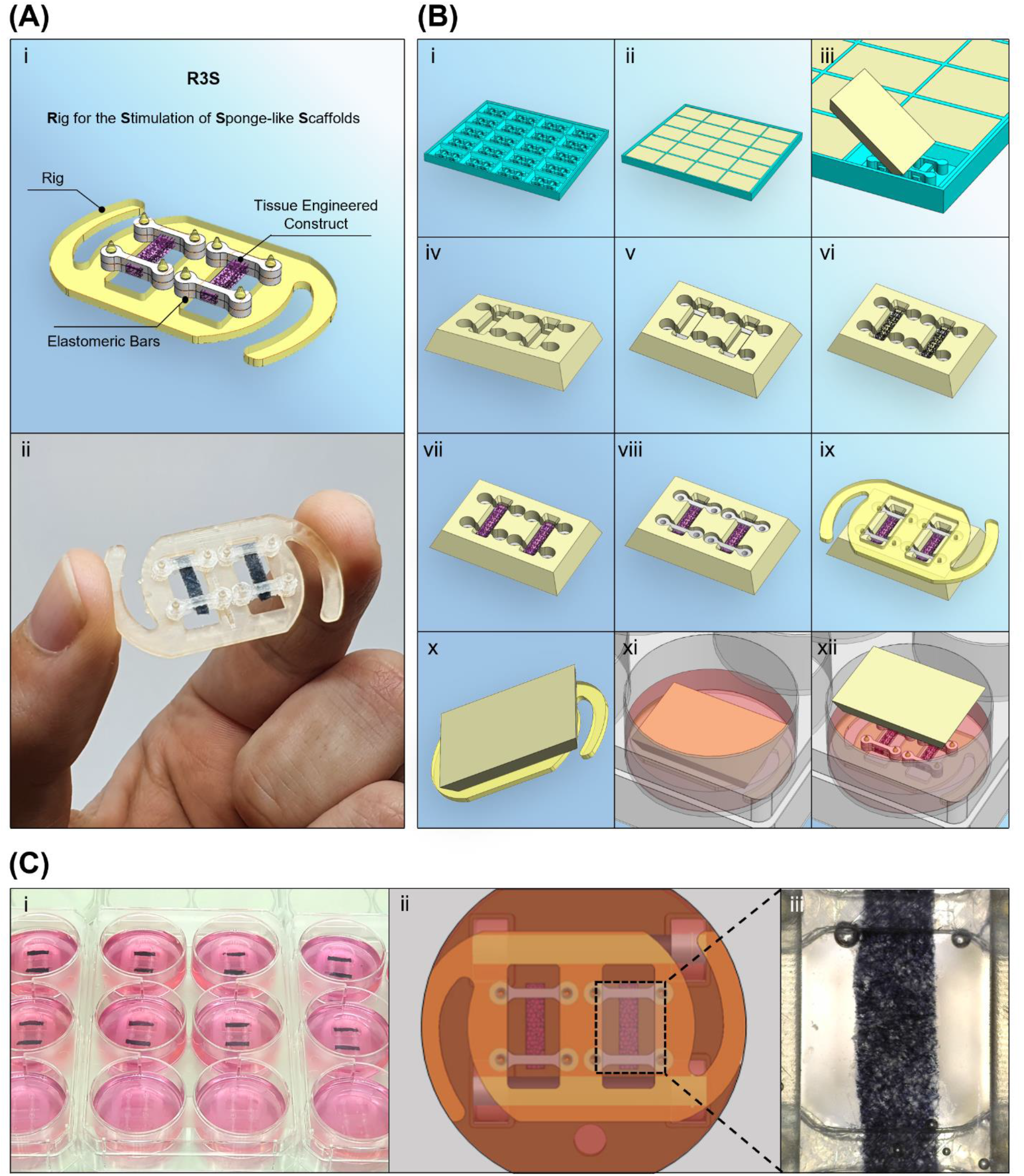
**(A)** (i) Schematic and (ii) picture of R3S (Rig for the Stimulation of Sponge-like Scaffolds). **(B)** (i-iv) Sequence of preparation of seeding chambers (v-xii) Sequence of assembly of R3S. **(C)** Assembled cell culture setup, showing (i) a picture of a set of plates with R3S, (ii) a schematic of the view from the bottom of the plate with focus on a single well and finally (iii) a picture obtained with the microscope camera focusing on a single scaffold.

### 2.6. Cell culture and evaluation of biocompatibility

C3H10 mouse embryonic fibroblasts (ATCC® CCL-226TM) were cultured in growth media prepared with Dulbecco’s Modified Eagle’s Medium (DMEM) low glucose (Sigma-Aldrich) containing 10 v/v% foetal bovine serum (FBS) (Gibco® by Life Technologies) and 2 v/v% Penicillin Streptomycin (Pen-Strep) (Sigma Aldrich) at 37 °C with 5 v/v% CO2. Cells between passages 12 and 15 were used. 3D scaffolds were sterilised immediately prior to use undergoing multiple cycles of incubation in 70 v/v% ethanol under UV light, followed by sterile deionised water, and finally incubated overnight in growth media for 24 hours.

### 2.7. Seeding protocol and validation of pacing bioreactor with C3H10

To facilitate accurate assembly of the rig and for consistency between replicates, while also confining the cell seeding volume, specific seeding chambers were designed. These were obtained from the addition of sterile 2 w/v% agarose solution into a 3D printed mould specifically designed to fit rectangular sponge-like scaffolds with dimensions of 2 × 1 × 9 mm and four elastomeric bars per scaffold (**Figure 3.A**, **S5** and **Video 4**). The agarose solution solidified in a few minutes, then making it possible to remove the seeding chamber and proceed with the in vitro experiment.

For validation of the pacing bioreactor and of R3S, C3H10 were seeded as illustrated in **Figure 3.B** and **Video 5**. Specifically, two elastomeric bars were set in each compartment of the seeding chamber. Scaffolds were added, and any excess liquid removed. The cell suspension was mixed in equal amount with Matrigel™ to reach a final seeding volume of 20 μl and 250k cells that was pipetted onto the dry scaffold. After an incubation of 30 minutes at 37 °C, to allow the transition from liquid to gel of the media/Matrigel, two more elastomeric bars were placed on top of the scaffold in a sandwich-like fashion. At this stage, the main body of R3S was pressed into the seeding chamber, piercing the agarose and locking in position the elastomeric bars and the scaffolds. The setup could be flipped and moved to a 6-well plate which was prefilled with culture media. After one more hour of incubation, the agarose seeding chamber was removed (**Figure 3.C** and **Video 6)**. Starting from day 3, cells were paced with a biphasic pulsatile regime in the order of ±2.5 V, 2 ms pulse, 2 Hz. A daily 1 hour long conditioning session was carried out until day 7 (**Video 7**).

### 2.8. Live/dead assay

Cell viability was assessed using a live/dead assay implementing a solution of 2 μl/ml Ethidium Homodimer and 0.5 μl/ml Calcein (Cambridge Bioscience) in PBS. This solution was incubated at 37°C for 1 hour and afterwards washed three times with PBS. Samples were kept at 37°C until imaged. Imaging was performed with a Leica SP8 scanning confocal microscope (Leica Microsystems, Germany). For quantification, a minimum of 3 pictures per experimental replicate were taken and subsequently analysed with ImageJ. Pictures were obtained as resultant z-projection of z-stack with different z-step and depth of penetration from the surface. Specifically, z-step size of 25 μm and depths of 125 μm were used. Cell viability was defined as the ratio of live cells over the total cell number (%), while the live cell density was obtained as ratio between live and total cells normalised by the ROI area (cells/mm^2^).

### 2.9. DAPI/Phalloidin fluorescent staining

Cell spreading was evaluated using cytofluorescent staining. After three washing in PBS, samples were fixed in 4 w/v% paraformaldehyde for 60 minutes at room temperature. Following three more washings in PBS, samples were incubated within a working dye solution prepared with 1 μl/ml phalloidin (Santa Cruz Technology, USA) and 4’,6-Diamidine-2’-phenylindoledihydrochloride (DAPI, 1 mg/ml, Sigma-Aldrich, Ireland), to highlight filamentous actin (f-actin) of the cell cytoskeleton and cell nuclei respectively. Micrographs were obtained using either Leica SP8 scanning confocal microscope (Leica Microsystems, Germany), and image processing for nuclei orientation was performed with custom-made scripts in Matlab®.

### 2.10. AlamarBlue™ metabolic assay

To evaluate cell condition, a standard AlamarBlue™ metabolic assay was performed. After removal of cell culture media and one washing with PBS, a AlamarBlue™ working solution containing 20 of AlamarBlue™ reagent was added to the culture and incubated for 1 hour. Media was gently mixed with pipette and incubated for 1 hour; afterwards the media was mixed again and moved to 96-well plate for analysis.

### 2.11. Picogreen™ assay

Biochemical assay was used to identify DNA content using a Picogreen™ assay. At specific timepoints, samples were washed in PBS and frozen at −80°C until the moment of the assay when a 18 hours digestion in papain enzymatic solution took place. Analysis was then performed accordingly to the provided standard protocol.

### 2.12. Statistical analysis

Statistical analysis was performed using GraphPad Prism 9 (GraphPad Software, USA). Where appropriate a one-way or two-way analysis of variance (ANOVA) followed by Tukey’s multiple comparison. If not otherwise specified, results are presented as mean ± standard deviation and differences are considered as statistically significant for p < 0.05.

## 3. Results and Discussion

In this study, we describe a setup for securing and pacing 3D sponge-like scaffolds, with a custom-made bioreactor to provide electrical pacing; providing detailed drawing and assembly videos to achieve this. To promote the adhesion of cultured cells to the scaffolds and implement a level of standardisation during assembly of the R3S system, a detailed seeding protocol was optimised. The seeding chambers were fabricated with agarose, and cells were suspended and delivered within a weak Matrigel™ mix (1:1 ratio mixed with cell suspension in media) which undergoes a rapid sol/gel transition. The rationale for this seeding is that the assembly of R3S involves a compression manoeuvre by the elastomeric bars on the scaffolds, as well as permanent suspension of the scaffolds in a floating, stable position at constant distance from the bottom of the plate. These factors may hinder cell delivery to the scaffolds and subsequent seeding density. At day 1, DNA content between groups seeded with C3H10 with standard media in a PDMS mould or in R3S was comparable, and no statistical difference was found between the different scaffold types, confirming that the use of the Matrigel™ mix (1:1) led to a successful seeding yield (**Figure S6**). Samples seeded with C3H10 in R3S were then studied up to seven days, applying electrical pacing with this in-house bioreactor and their performance compared to day 3. Electrical cues have the potential to ameliorate cell performance and influence differentiation fate, however, the application of an improper electrical signal can drive uneven electric field distribution, excess of ions with generation of by-products, all eventually leading to detrimental effect on cell viability ^30^. Pacing patterns, pulse duration, peak intensity, pulse frequency, and duration of the pacing session are all key players in the effective delivery of pacing ^31^. In particular, the adoption of a biphasic pulse instead of monophasic has shown to reduce the risk of by-products production by limiting the formation of nonreversible-Faradaic reactions ^12^. The choice of a biphasic pulsatile stimulation and 1-hour long training sessions have been implemented with a conservative approach to preserve cell wellbeing and prevent excessive production of potentially harmful redox oxygen species. The findings presented here are proof that such choices successfully maintained cells viability and provides sufficient signalling cues to obtain a positive effect on their activity As demonstrated by quantification of live/dead imaging, fibroblasts remained viable at both day 3 and 7, with no difference between collagen alone, or PEDOT:PSS-based sponges, with or without electrical pacing (**Figure 4**). From the mosaic micrographs, cells were clearly distributed homogeneously across the scaffolds, with the exception of the terminal regions, corresponding to the sites of anchorage with the elastomeric bars. Here, the elastomeric bars held the scaffold ends without breaking them, and by day 7 a plastic compression of the scaffolds occurred together with less cell presence. The use of elastomeric bars is meant to not only simplify the assembly sequence, but also to maintain scaffolds in a free-floating location in the media. Moreover, acting as flexible constrains with known stiffness, these bars are particularly important for application of this rig with mechanically dynamic and contracting cells such as those from the cardiac or skeletal muscles. Indeed, monitoring and quantification of the beating of these cells are a crucial towards the understanding of their physiology. Real-time monitoring during the experiment will enables the indirect quantification of the in vitro models, for example the frequency of contraction or derivatisation twitch forces generated ^32^ (**Figure 1B-iii**). DNA quantification verified cell proliferation within all groups, with trends of increased proliferation for paced groups when compared with controls (**Figure 5.A**). Similar results were obtained for the analysis of cell metabolism; in particular, cells on paced anisotropic scaffolds demonstrated significant increases in metabolism compared to all collagen and isotropic groups, suggesting a benefit on cell performance by the synergistic effect of an anisotropic morphology and the external pacing (**Figure 5.B**).

**Figure 4.**
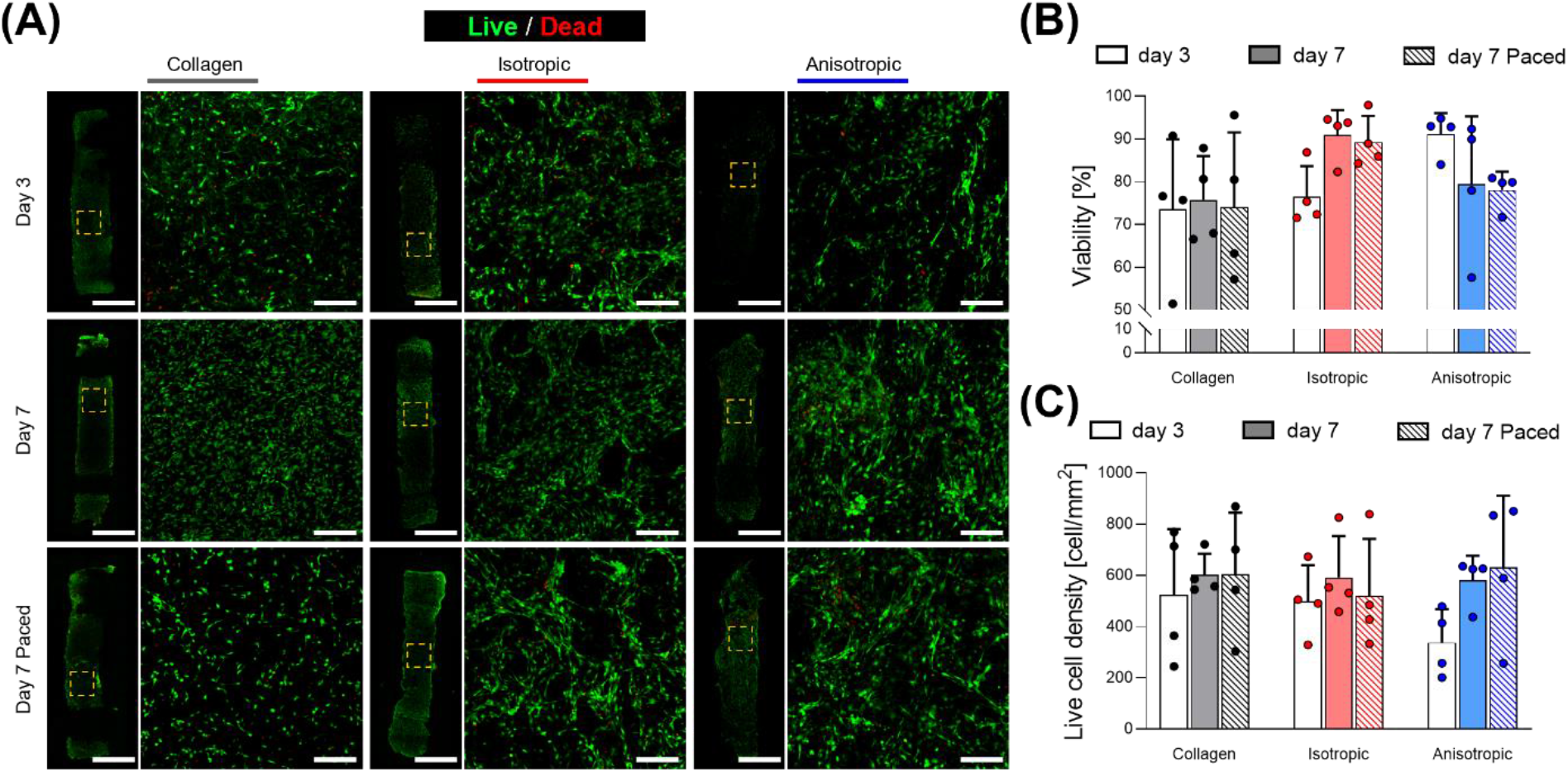
Effects of pacing C3H10 cells in scaffolds on viability. **(A)** Micrographs from confocal microscope fluorescent staining for live/dead of CH310 cells at days 3, 7 with and without pacing. **(B, C)** Quantification of viability (extracted from live/dead staining) at days 3, 7 with and without pacing quantified as viability and alive cell density (n=4). Scale bars: A = 200 μm; A insets = 200 μm. Bar graphs demonstrate the mean with error bars representing standard deviation. Data values are presented as associated points. Statistical significance was performed using two-way ANOVA with Tukey’s post-hoc test.

**Figure 5.**
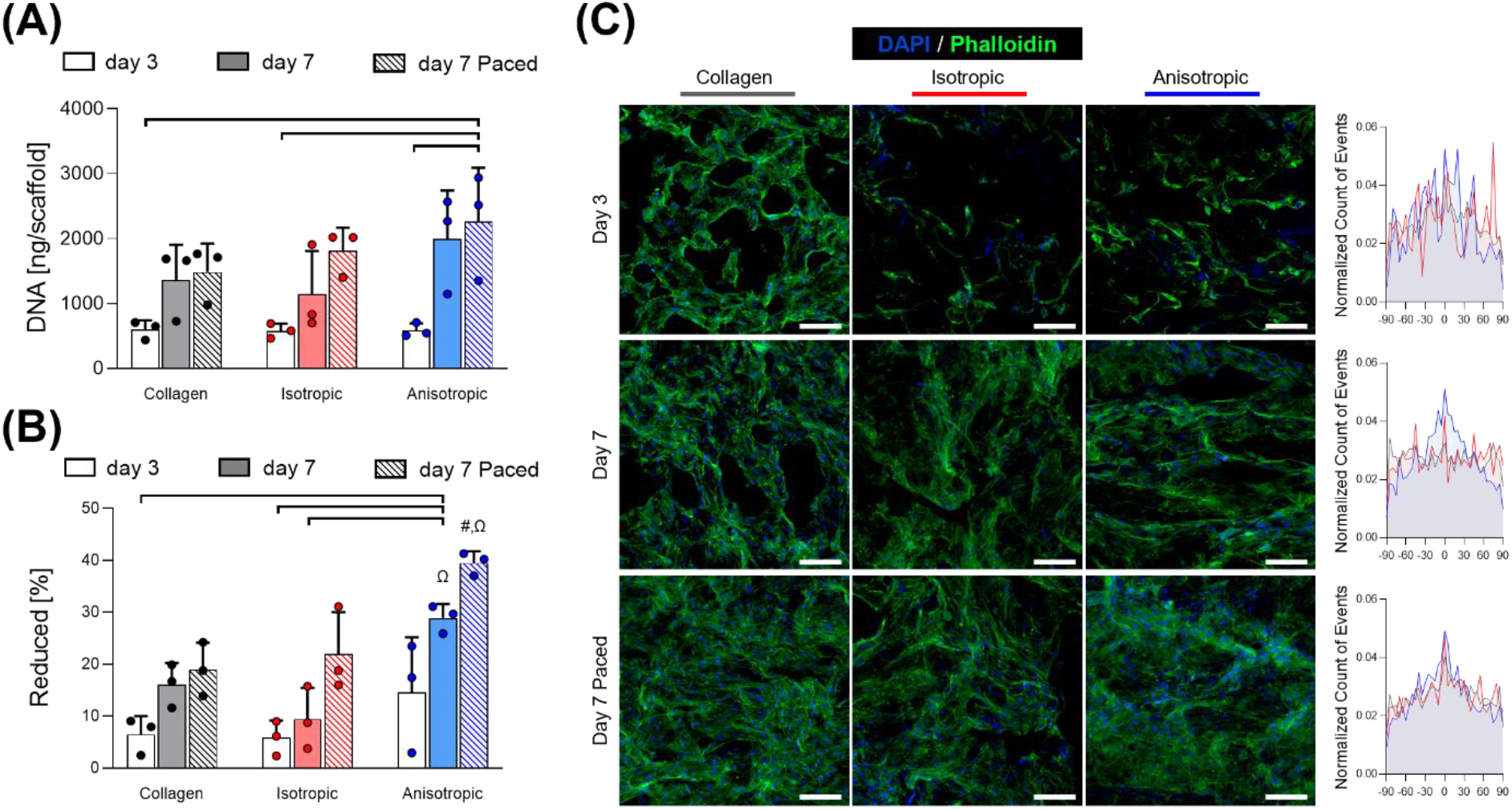
Effects of pacing C3H10 cells in scaffolds on cell proliferation, metabolism and orientation. **(A)** Quantification of DNA via Picogreen™ assay, expressed as ng per scaffold (n=3). **(B)** AlamarBlue™ assay performed on scaffolds (n=3). **(C)** Micrographs from confocal microscope fluorescent staining for nuclei/f-actin (DAPI and Phalloidin respectively) and distribution of nuclei orientation. Scale bars: C = 100 μm. Bar graphs demonstrate the mean with error bars representing standard deviation. Data values are presented as associated points. * and # represent statistical significance (p<0.05) (* between indicated groups, # with all other groups), while Ω is shared by groups that are not statistically significant using two-way ANOVA with Tukey’s post-hoc test.

In terms of cell morphology, no major qualitative difference was observed; when focussing on cell orientation as it related to nuclei orientation, it could be observed that cells aligned along the lamellae of the anisotropic scaffolds already from day 3 and they maintained such directionality up to day 7. As anticipated, cells on collagen or isotropic substrates did not have any preferential alignment at either day 3 or at day 7 without pacing, however this alignment adopted preferentially in parallel to the electric field of those samples that were subject to electrical pacing. While more studies are necessary to further validate and finalise the present results, here it is confirmed that orientation of C3H10 cell line can be influenced by either scaffold geometry or external pacing and indeed this bioreactor system facilitates this, similar to other cell types such as neural stem cells PC-12 ^33^ and H9C2 rat cardiomyoblast ^34^. Such is alignment hypothesized to be based on mechanisms that include voltage-gated ion channels, G-protein coupling receptors, integrins, cell polarization, and endogenous electric fields ^35–37^.

## 4. Conclusions

In this work we explain the design, fabrication, and validation of an electrical pacing bioreactor and of a Rig for Stimulation of Sponge-like Scaffolds (R3S) which allows for real time monitoring of these such systems. These can be manufactured inexpensively and in a scalable manner using rapid prototyping. The development of electroconductive scaffolds and electric-field stimulation models in vitro model is becoming increasingly pursued by researchers, there is a need for systems that can combine these two elements and often on that is easy (and cheap) to fabricate.

This system is not just limited to use with 3D porous scaffolds, and the electrical pacing bioreactor can be used to pace monolayer cultures, and the optical accessibility of the system enables an assessment of chemotaxis or electrotaxis ^38^. We presented an assembly sequence that is suitable for scaffolds generated with multiple material types and potentially different manufacturing techniques. The impact and validation of this platform is clear. When applying external pacing to C3H10 cells no detrimental effect on cell viability was observed, while a synergistic increase in metabolism and cell orientation was found with the combination of anisotropic scaffolds and pacing. Pacing was instrumental in cell alignment whether platforms were conductive, anisotropically aligned; or not. This system opens up great opportunity to study the interplay between cells and a wider repertoire of biomaterials that are prefabricated and predesigned, while observing the movement and contraction of these cellularised tissues in vitro.

## Supporting information

Video 1 - Fabrication of electrodes

Video 2 - Assembly of bioreactor

Video 3 - R3S overview

Video 4 - Fabrication of seeding chamber

Video 5 - Assembly of scaffolds into chambers and seeding

Video 6 - Coupling R3S with seeding chamber

Video 7 - Final setup

## ASSOCIATED CONTENT

The following files are available free of charge.

### Supporting Information (PDF)

Figure S1: Drawings of the components of the mould for the embedding of PDMS on carbon bars for the development of electrodes.

Figure S2: Drawing of the lid adapter of the bioreactor.

Figure S3: Electrical circuit schematic of the Pacing Bioreactor.

Figure S4: Drawing of R3S.

Figure S5: Drawings of Mould for Seeding chambers and of a single seeding chamber.

Figure S6: Effects of seeding technique of C3H10 cells on scaffolds on cell proliferation.

### Videos (MP4)

Video 1 - Fabrication of electrodes

Video 2 - Assembly of bioreactor

Video 3 - R3S overview

Video 4 - Fabrication of seeding chamber

Video 5 - Assembly of scaffolds into chambers and seeding

Video 6 - Coupling R3S with seeding chamber

Video 7 - Final setup

## AUTHOR INFORMATION

### Authors

Matteo Solazzo

Department of Mechanical, Manufacturing and Biomedical Engineering, Trinity Biomedical Sciences Institute, Trinity College Dublin, 152–160 Pearse Street, Dublin 2, Ireland.

## ACKNOWLEDGMENT

This work was supported through an SFI-HRB Wellcome Trust-ISSF Award Institutional Strategic Support Fund (Ref No. 204814/Z/16/Z), the Irish Research Council (Project No. GOIPG/2019/818) and Science Foundation Ireland (SFI), Ireland, through the Advanced Materials and Bioengineering Research (AMBER) Centre (SFI/12/RC/2278_P2, partly supported by the European Regional Development Fund).

## ABBREVIATIONS

3D: three-dimensional
DAPI: 4’,6-Diamidine-2’-phenylindoledihydrochloride
EDC: *N*-(*3*-Dimethylaminoproypl)-*N*-ethylcarbodiimide hydrochloride
GOPS: glycidoxipropyl-trimethoxysilane
NHS: *N*-Hydroxysuccinimide
PDMS: polydimethylsiloxane
PEDOT:PSS: poly(3,4-ethylenedioxythiophene):polystyrene sulfonate
R3S: rig for stimulation of sponge-like scaffolds
ROI: region of interest

## REFERENCES

1. Martin, I.; Wendt, D.; Heberer, M., The role of bioreactors in tissue engineering. Trends in Biotechnology 2004, 22 (2), 80–86.

2. Zhao, Y.; Rafatian, N.; Feric, N. T.; Cox, B. J.; Aschar-Sobbi, R.; Wang, E. Y.; Aggarwal, P.; Zhang, B.; Conant, G.; Ronaldson-Bouchard, K.; Pahnke, A.; Protze, S.; Lee, J. H.; Davenport Huyer, L.; Jekic, D.; Wickeler, A.; Naguib, H. E.; Keller, G. M.; Vunjak-Novakovic, G.; Broeckel, U.; Backx, P. H.; Radisic, M., A Platform for Generation of Chamber-Specific Cardiac Tissues and Disease Modeling. Cell 2019, 176 (4), 913–927 e18.

3. Latchoumane, C.-F. V.; Jackson, L.; Sendi, M. S. E.; Tehrani, K. F.; Mortensen, L. J.; Stice, S. L.; Ghovanloo, M.; Karumbaiah, L., Chronic Electrical Stimulation Promotes the Excitability and Plasticity of ESC-derived Neurons following Glutamate-induced Inhibition In vitro. Scientific Reports 2018, 8 (1), 10957.

4. Nikolić, N.; Skaret Bakke, S.; Tranheim Kase, E.; Rudberg, I.; Flo Halle, I.; Rustan, A. C.; Thoresen, G. H.; Aas, V., Electrical Pulse Stimulation of Cultured Human Skeletal Muscle Cells as an In Vitro Model of Exercise. PLOS ONE 2012, 7 (3), e33203.

5. Sebastian, A.; Volk, S. W.; Halai, P.; Colthurst, J.; Paus, R.; Bayat, A., Enhanced Neurogenic Biomarker Expression and Reinnervation in Human Acute Skin Wounds Treated by Electrical Stimulation. J Invest Dermatol 2017, 137 (3), 737–747.

6. Su, C. Y.; Fang, T.; Fang, H. W., Effects of Electrostatic Field on Osteoblast Cells for Bone Regeneration Applications. Biomed Res Int 2017, 2017, 7124817.

7. Gu, J.; He, X.; Chen, X.; Dong, L.; Weng, W.; Cheng, K., Effects of electrical stimulation on cytokine-induced macrophage polarization. J Tissue Eng Regen Med 2022, 16 (5), 448–459.

8. Weinberger, F.; Breckwoldt, K.; Pecha, S.; Kelly, A.; Geertz, B.; Starbatty, J.; Yorgan, T.; Cheng, K. H.; Lessmann, K.; Stolen, T.; Scherrer-Crosbie, M.; Smith, G.; Reichenspurner, H.; Hansen, A.; Eschenhagen, T., Cardiac repair in guinea pigs with human engineered heart tissue from induced pluripotent stem cells. Sci Transl Med 2016, 8 (363), 363ra148.

9. Li, J.; Zhang, L.; Yu, L.; Minami, I.; Miyagawa, S.; Horning, M.; Dong, J.; Qiao, J.; Qu, X.; Hua, Y.; Fujimoto, N.; Shiba, Y.; Zhao, Y.; Tang, F.; Chen, Y.; Sawa, Y.; Tang, C.; Liu, L., Circulating re-entrant waves promote maturation of hiPSC-derived cardiomyocytes in self-organized tissue ring. Commun Biol 2020, 3 (1), 122.

10. Querdel, E.; Reinsch, M.; Castro, L.; Kose, D.; Bahr, A.; Reich, S.; Geertz, B.; Ulmer, B.; Schulze, M.; Lemoine, M. D.; Krause, T.; Lemme, M.; Sani, J.; Shibamiya, A.; Studemann, T.; Kohne, M.; Bibra, C. V.; Hornaschewitz, N.; Pecha, S.; Nejahsie, Y.; Mannhardt, I.; Christ, T.; Reichenspurner, H.; Hansen, A.; Klymiuk, N.; Krane, M.; Kupatt, C.; Eschenhagen, T.; Weinberger, F., Human Engineered Heart Tissue Patches Remuscularize the Injured Heart in a Dose-Dependent Manner. Circulation 2021, 143 (20), 1991–2006.

11. Eschenhagen, T.; Fink, C.; Remmers, U.; Scholz, H.; Wattchow, J.; Weil, J.; Zimmermann, W.; Dohmen, H. H.; Schafer, H.; Bishopric, N.; Wakatsuki, T.; Elson, E. L., Three-dimensional reconstitution of embryonic cardiomyocytes in a collagen matrix: a new heart muscle model system. FASEB journal : official publication of the Federation of American Societies for Experimental Biology 1997, 11 (8), 683–94.

12. Tandon, N.; Cannizzaro, C.; Chao, P. H.; Maidhof, R.; Marsano, A.; Au, H. T.; Radisic, M.; Vunjak-Novakovic, G., Electrical stimulation systems for cardiac tissue engineering. Nat Protoc 2009, 4 (2), 155–73.

13. Balint, R.; Cassidy, N. J.; Cartmell, S. H., Electrical stimulation: a novel tool for tissue engineering. Tissue engineering. Part B, Reviews 2013, 19 (1), 48–57.

14. Murphy, J. F.; Mayourian, J.; Stillitano, F.; Munawar, S.; Broughton, K. M.; Agullo-Pascual, E.; Sussman, M. A.; Hajjar, R. J.; Costa, K. D.; Turnbull, I. C., Adult human cardiac stem cell supplementation effectively increases contractile function and maturation in human engineered cardiac tissues. Stem Cell Res Ther 2019, 10 (1), 373.

15. Turnbull, I. C.; Mayourian, J.; Murphy, J. F.; Stillitano, F.; Ceholski, D. K.; Costa, K. D., Cardiac Tissue Engineering Models of Inherited and Acquired Cardiomyopathies. Methods Mol Biol 2018, 1816, 145–159.

16. Freed, L. E.; Vunjak-Novakovic, G., Microgravity tissue engineering. In Vitro Cell Dev Biol Anim 1997, 33 (5), 381–5.

17. Radisic, M.; Euloth, M.; Yang, L.; Langer, R.; Freed, L. E.; Vunjak-Novakovic, G., High-density seeding of myocyte cells for cardiac tissue engineering. Biotechnol Bioeng 2003, 82 (4), 403–14.

18. Zhou, J.; Chen, J.; Sun, H.; Qiu, X.; Mou, Y.; Liu, Z.; Zhao, Y.; Li, X.; Han, Y.; Duan, C.; Tang, R.; Wang, C.; Zhong, W.; Liu, J.; Luo, Y.; Mengqiu Xing, M.; Wang, C., Engineering the heart: evaluation of conductive nanomaterials for improving implant integration and cardiac function. Sci Rep 2014, 4, 3733.

19. Chen, S.; Hsieh, M. H.; Li, S. H.; Wu, J.; Weisel, R. D.; Chang, Y.; Sung, H. W.; Li, R. K., A conductive cell-delivery construct as a bioengineered patch that can improve electrical propagation and synchronize cardiomyocyte contraction for heart repair. J Control Release 2020, 320, 73–82.

20. Wang, L.; Jiang, J.; Hua, W.; Darabi, A.; Song, X.; Song, C.; Zhong, W.; Xing, M. M. Q.; Qiu, X., Mussel-Inspired Conductive Cryogel as Cardiac Tissue Patch to Repair Myocardial Infarction by Migration of Conductive Nanoparticles. Advanced Functional Materials 2016, 26 (24), 4293–4305.

21. Balint, R.; Cassidy, N. J.; Cartmell, S. H., Conductive polymers: towards a smart biomaterial for tissue engineering. Acta Biomater 2014, 10 (6), 2341–53.

22. Ryan, A. J.; Kearney, C. J.; Shen, N.; Khan, U.; Kelly, A. G.; Probst, C.; Brauchle, E.; Biccai, S.; Garciarena, C. D.; Vega-Mayoral, V.; Loskill, P.; Kerrigan, S. W.; Kelly, D. J.; Schenke-Layland, K.; Coleman, J. N.; O’Brien, F. J., Electroconductive Biohybrid Collagen/Pristine Graphene Composite Biomaterials with Enhanced Biological Activity. Adv Mater 2018, 30 (15), e1706442.

23. Bray, M. A.; Sheehy, S. P.; Parker, K. K., Sarcomere alignment is regulated by myocyte shape. Cell Motil Cytoskeleton 2008, 65 (8), 641–51.

24. Wilson, A. J.; Schoenauer, R.; Ehler, E.; Agarkova, I.; Bennett, P. M., Cardiomyocyte growth and sarcomerogenesis at the intercalated disc. Cell Mol Life Sci 2014, 71 (1), 165–81.

25. Engler, A. J.; Sen, S.; Sweeney, H. L.; Discher, D. E., Matrix elasticity directs stem cell lineage specification. Cell 2006, 126 (4), 677–89.

26. Solazzo, M.; Monaghan, M. G., Structural crystallisation of crosslinked 3D PEDOT:PSS anisotropic porous biomaterials to generate highly conductive platforms for tissue engineering applications. Biomater. Sci. 2021, 9 (12), 4317–4328.

27. Solazzo, M.; Hartzell, L.; O’Farrell, C.; Monaghan, M. G., Beyond Chemistry: Tailoring Stiffness and Microarchitecture to Engineer Highly Sensitive Biphasic Elastomeric Piezoresistive Sensors. ACS Appl Mater Interfaces 2022, 14 (17), 19265–19277.

28. Yang, C., Enhanced physicochemical properties of collagen by using EDC/NHS-crosslinking. Bulletin of Materials Science 2012, 35 (5), 913–918.

29. Hansen, A.; Eder, A.; Bonstrup, M.; Flato, M.; Mewe, M.; Schaaf, S.; Aksehirlioglu, B.; Schwoerer, A. P.; Uebeler, J.; Eschenhagen, T., Development of a drug screening platform based on engineered heart tissue. Circ Res 2010, 107 (1), 35–44.

30. Nuccitelli, R.; Lui, K.; Kreis, M.; Athos, B.; Nuccitelli, P., Nanosecond pulsed electric field stimulation of reactive oxygen species in human pancreatic cancer cells is Ca(2+)-dependent. Biochem Biophys Res Commun 2013, 435 (4), 580–5.

31. Lou, L.; Lopez, K. O.; Nautiyal, P.; Agarwal, A., Integrated Perspective of Scaffold Designing and Multiscale Mechanics in Cardiac Bioengineering. Advanced NanoBiomed Research 2021, 1 (12), 2100075.

32. Rodriguez, M. L.; Graham, B. T.; Pabon, L. M.; Han, S. J.; Murry, C. E.; Sniadecki, N. J., Measuring the contractile forces of human induced pluripotent stem cell-derived cardiomyocytes with arrays of microposts. J Biomech Eng 2014, 136 (5), 051005.

33. Chen, C.; Ruan, S.; Bai, X.; Lin, C.; Xie, C.; Lee, I. S., Patterned iridium oxide film as neural electrode interface: Biocompatibility and improved neurite outgrowth with electrical stimulation. Mater Sci Eng C Mater Biol Appl 2019, 103, 109865.

34. Ganji, Y.; Li, Q.; Quabius, E. S.; Bottner, M.; Selhuber-Unkel, C.; Kasra, M., Cardiomyocyte behavior on biodegradable polyurethane/gold nanocomposite scaffolds under electrical stimulation. Mater Sci Eng C Mater Biol Appl 2016, 59, 10–18.

35. Li, Y.; Huang, G.; Zhang, X.; Wang, L.; Du, Y.; Lu, T. J.; Xu, F., Engineering cell alignment in vitro. Biotechnology Advances 2014, 32 (2), 347–365.

36. Banks, T. A.; Luckman, P. S. B.; Frith, J. E.; Cooper-White, J. J., Effects of electric fields on human mesenchymal stem cell behaviour and morphology using a novel multichannel device. Integrative Biology 2015, 7 (6), 693–712.

37. Lang, M.; Bunn, S.; Gopalakrishnan, B.; Li, J., Use of weak DC electric fields to rapidly align mammalian cells. Journal of Neural Engineering 2021, 18 (5), 054002.

38. Lin, F.; Baldessari, F.; Gyenge, C. C.; Sato, T.; Chambers, R. D.; Santiago, J. G.; Butcher, E. C., Lymphocyte electrotaxis in vitro and in vivo. J Immunol 2008, 181 (4), 2465–71.

