## Supporting Information for "A guide to the manufacture of sustainable, ready to use in vitro platforms for the electric-field pacing of cellularised 3D porous scaffolds"

* Prof. M. G. Monaghan

Department of Mechanical, Manufacturing and Biomedical Engineering, Trinity Biomedical Sciences Institute, Trinity College Dublin, 152–160 Pearse Street, Dublin 2, Ireland.

1. Department of Mechanical, Manufacturing and Biomedical Engineering, Trinity College Dublin, Dublin 2, Ireland.
2. Advanced Materials and BioEngineering Research (AMBER) Centre at Trinity College Dublin and the Royal College of Surgeons in Ireland, Dublin 2, Ireland.
3. CÚRAM, Centre for Research in Medical Devices, National University of Ireland, Galway, Newcastle Road, H91 W2TY Galway, Ireland

Keywords: biomaterials, electroconductive biomaterials, cardiac tissue engineering, polymer chemistry, organoid generation

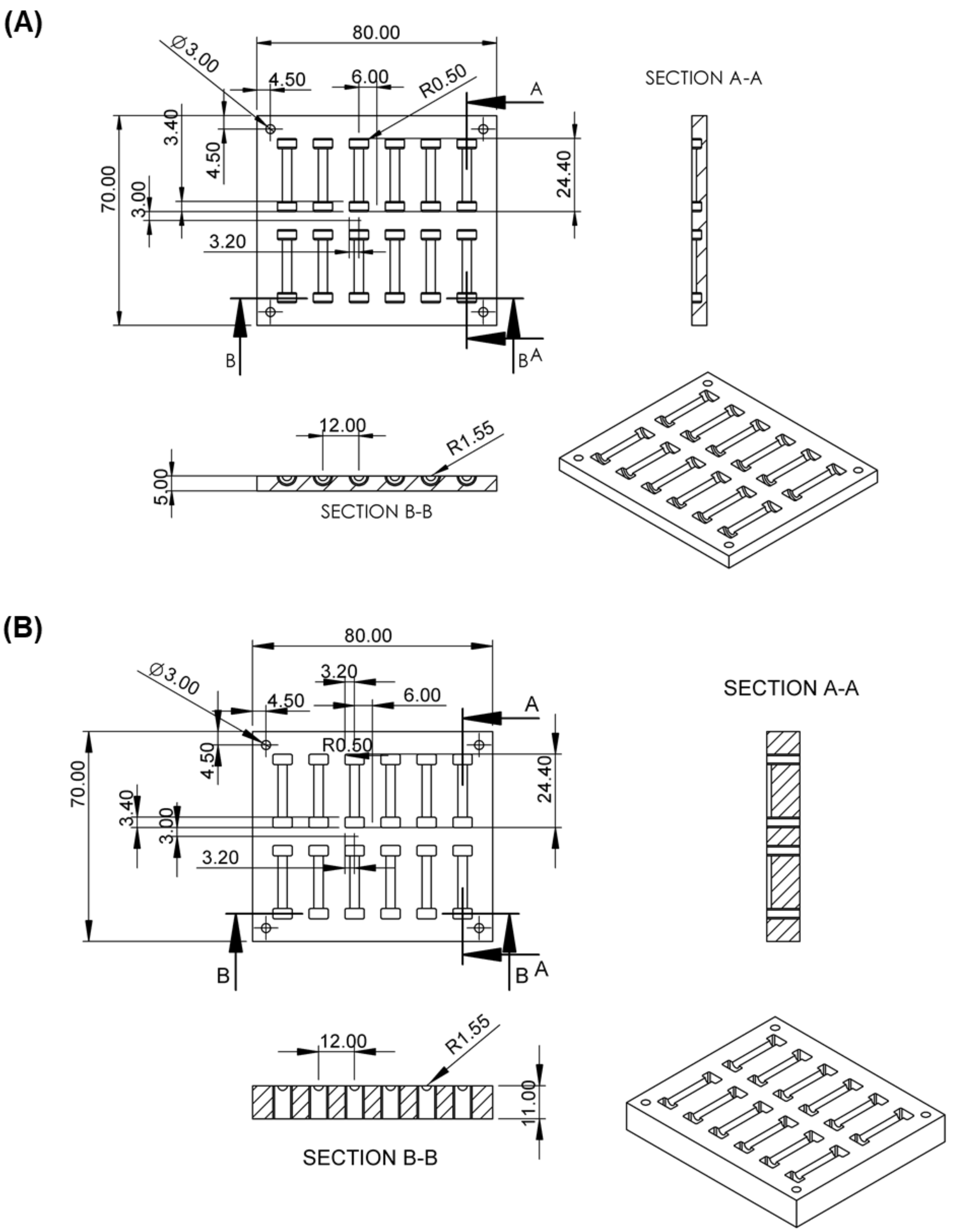


**Supporting Information 1.** Drawings of the Bottom **(A)** and Top **(B)** components of the mould for the embedding of PDMS on carbon bars for the development of electrodes. Dimensions are expressed in mm.

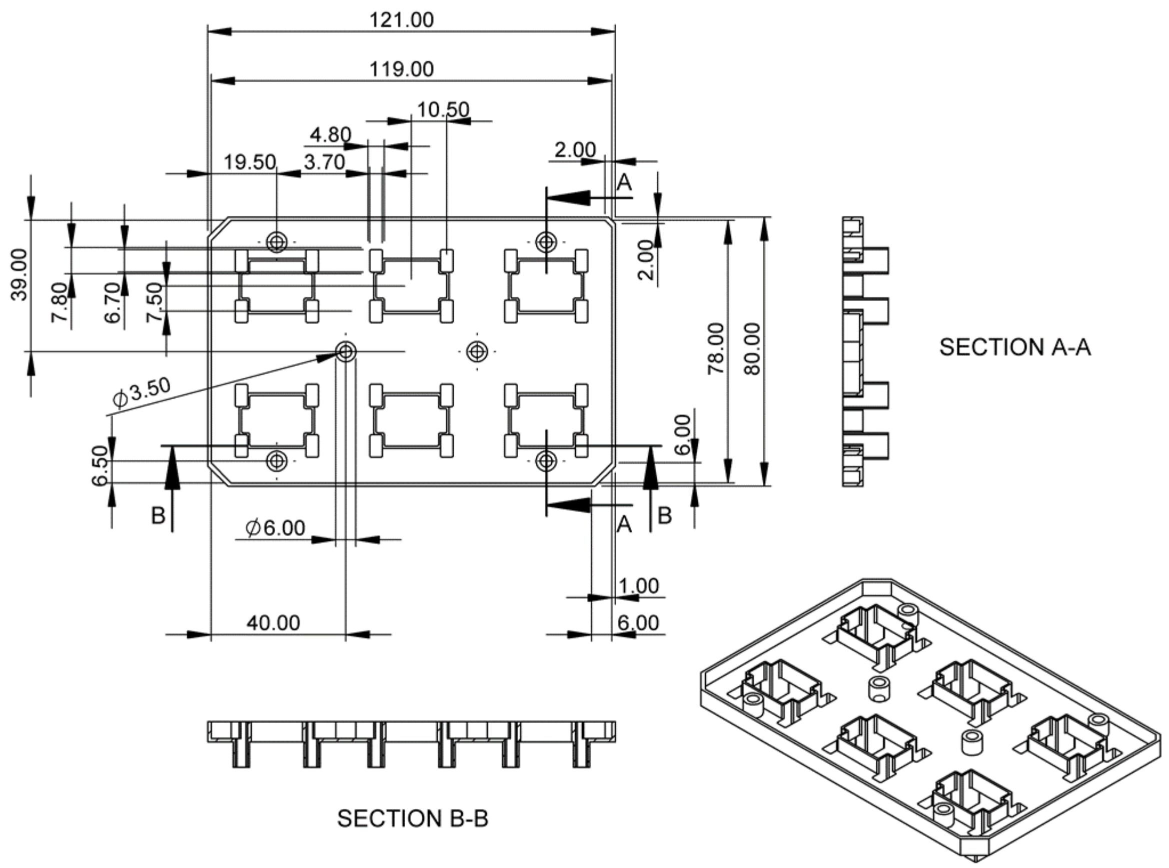


**Supporting Information 2**. Drawing of the lid adapter of the bioreactor. Dimensions are expressed in mm.

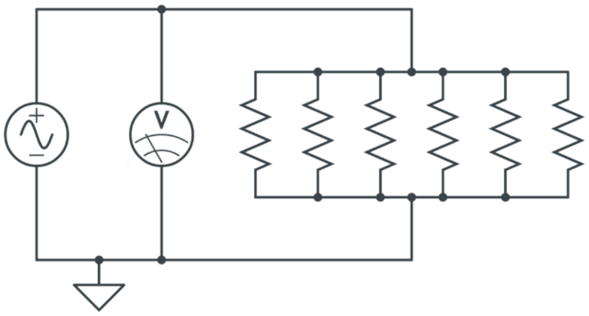


**Supporting Information 3**. Electrical circuit schematic of the Pacing Bioreactor

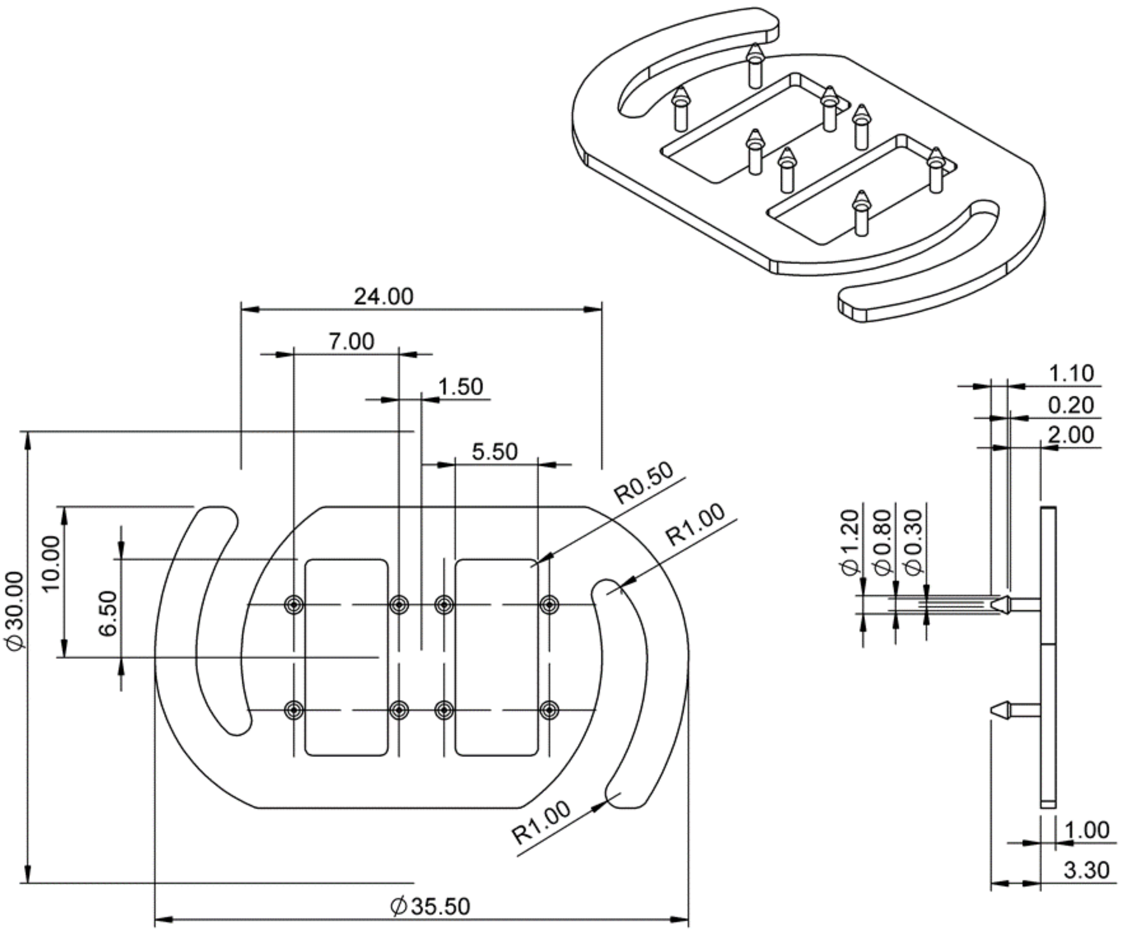


**Supporting Information 4**. Drawing of R3S - Rig for the Stimulation of Sponge-like Scaffolds. Dimensions are expressed in mm.

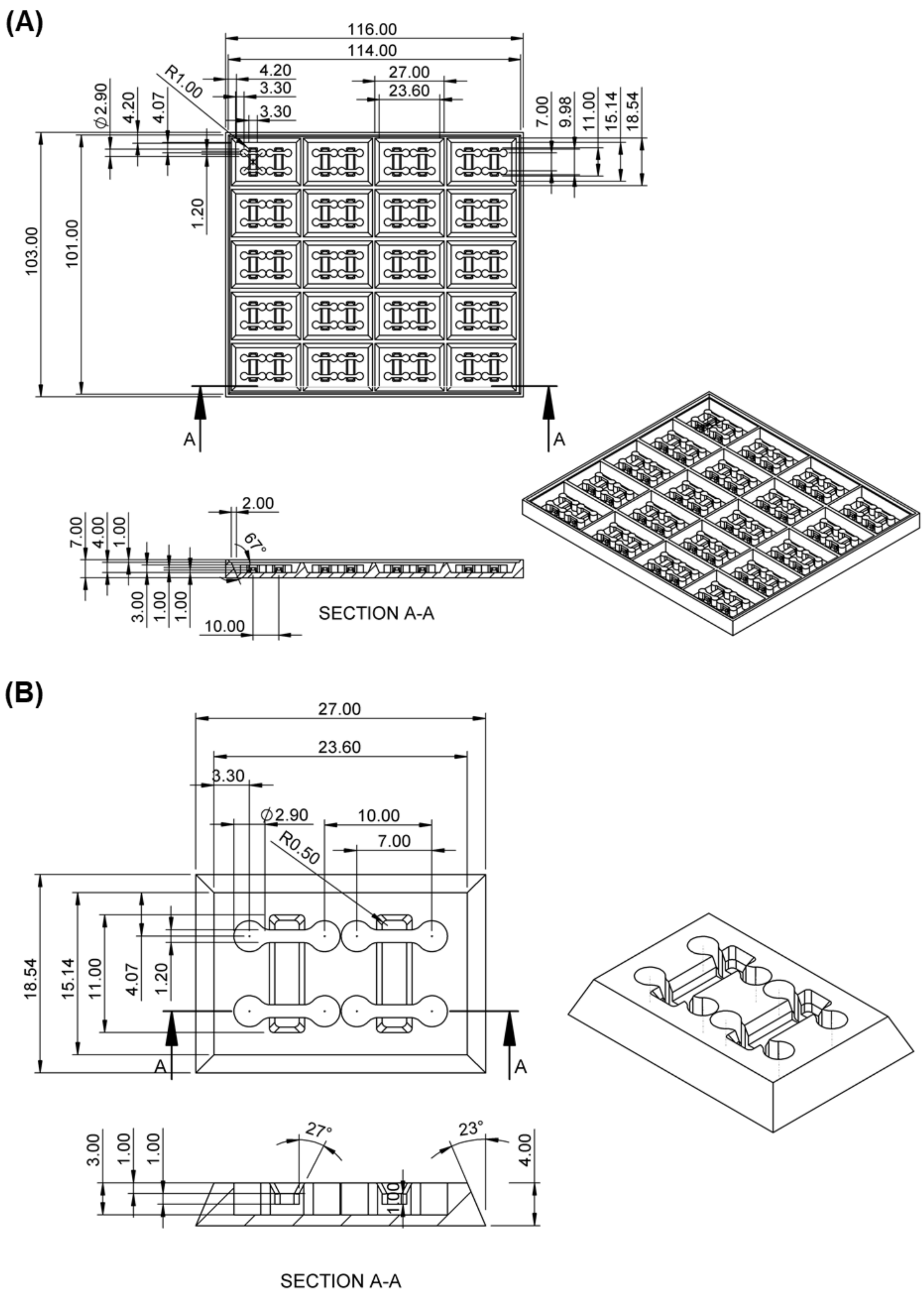


**Supporting Information 5.** Drawings of **(A)** Mould for Seeding chambers and of **(B)** a single seeding chamber. Dimensions are expressed in mm.

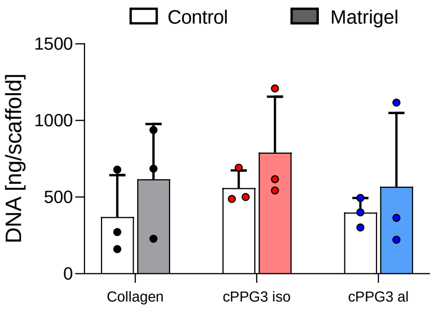


**Supporting Information 6**. Effects of seeding technique of C3H10 cells on scaffolds on cell proliferation. Quantification of DNA via Picogreen™ assay, expressed as ng per scaffold (n=3). Bar graphs demonstrate the mean with error bars representing standard deviation. Data values are presented as associated points. Statistically analysis was performed using two-way ANOVA with Tukey’s post-hoc test
